# The signature of reproduction on wood anatomy in European beech reveals potential for masting reconstruction

**DOI:** 10.1101/2025.05.27.656364

**Authors:** Giulia Resente, Alan Crivellaro, Emilie Fleurot, Alma Piermattei, Francesco Maimone, Martin Wilmking, Andrew Hacket-Pain, Renzo Motta, Davide Ascoli

**Affiliations:** Department of Agricultural, Forest and Food Sciences, University of Torino, Grugliasco (TO), Italy; Forest Biometrics Laboratory, Faculty of Forestry, "Stefan cel Mare" University of Suceava, Suceava, Romania; Institute of Botany and Landscape Ecology, University Greifswald, Greifswald, Germany; Department of Geography and Planning, School of Environmental Sciences, University of Liverpool, Liverpool, UK

**Author notes:** **Contact information** Giulia Resente.

**Keywords:** “mast seeding”, “dendroecology”, “climate change”, “quantitative wood anatomy”, “reconstruction”, “beech”, “reproduction”, “drought”

## Abstract

1. We investigated the potential of wood anatomical traits to improve the reconstruction of masting events—variable and synchronized patterns of seed production —which are key to understanding tree species’ responses to current and predicted climate variability. Traditional reliance on tree-ring width as a proxy for reproduction is limited, as growth reductions can also result from drought and other stressors.
2. We analyzed 12 beech cores from North-East Germany, building a 52-year dataset. A wide range of wood anatomical traits was assessed to disentangle the effects of masting and drought. We used multivariate regression and developed a random forest model to evaluate the predictive power of these traits compared to tree ring width alone.
3. Results suggested a complex mechanism of carbon reallocation towards reproduction, while reflecting a compensatory strategy to maintain hydraulic function and mechanical stability under resource-constrained conditions. Number of parenchyma cells, vessel density, and lignin content estimates emerged as key predictors for masting, outperforming tree-ring width in capturing the reproductive signal.
4. Our findings establish a novel link between wood anatomy and masting events, demonstrating that quantitative wood anatomical traits offer a more accurate and ecologically relevant approach for reconstructing past masting dynamics.

## Introduction

Masting, or mast seeding, refers to the reproduction strategy where plant populations exhibit highly variable and synchronized patterns of seed production (Pearse et al., 2021). While masting enhances reproductive fitness by increasing reproduction efficiency via economies of scale (Kelly and Sork, 2002), the associated demands of massive reproductive effort also affect plant growth and carbon stocks. These changes in resource allocation trigger major cascading ecological effects (Bogdziewicz et al., 2021) involving several ecosystem aspects, such as soil nutrient dynamics (Brumme et al., 2021), sporocarps fruiting reduction (Michaud et al., 2024), and shifts in animal population dynamics due to fluctuating food availability (Ostfeld et al., 2000).

Climate variation influences masting patterns on an inter-annual and decadal scale (Ascoli et al., 2021). Rising temperatures due to climate change have been influencing masting occurrence and changing masting patterns (Foest et al., 2024), with strong negative effects on seed production (Bogdziewicz et al., 2023). However, our understanding of the effects of the ongoing climate change on masting patterns is limited by the length of the available time series of masting, which are often too short for a comparison of their inter-annual variability with the fluctuations and decadal trends in climate. Consequently, the limited availability of long-term observations of masting prevents the evaluation of recent trends against long-term variability in masting (i.e., baseline variability) (Prebble et al., 2022; Wion et al., 2025). Additionally, available data is strongly biased geographically and taxonomically, with long-term records restricted to regions and species valued in forestry (Hacket-Pain et al., 2022).

Reconstructing past masting patterns is essential for understanding how plant reproductive behavior responds to long-term climate variability. By identifying how masting variability shifts across climatic phases, researchers can better anticipate the ecological consequences of ongoing climate change (Mundo et al., 2021; Prebble et al., 2022). In woody species such as Araucaria (*Araucaria araucana* (Molina) K. Koch), Oak (*Quercus robur* L.), and European beech (*Fagus sylvatica* L.), most reconstruction efforts to date have relied on tree-ring width as a proxy for reproductive investment (Drobyshev et al., 2014; Hacket-Pain et al., 2024; Koenig et al., 2020; Mundo et al., 2021). European beech in particular has emerged as a relevant species in masting research, with numerous studies demonstrating reduced growth during mast years, supporting the hypothesis of a direct growth–reproduction trade-off (Bajocco et al., 2021; Bogdziewicz et al., 2021; Drobyshev et al., 2014; Hacket-Pain et al., 2015; Nussbaumer et al., 2021; Stolz et al., 2023).

Despite the established use of growth rings for reconstruction purposes, the accuracy of dendrochronology remains limited by the challenge of disentangling the effects of masting from other radial growth-reducing factors such as drought, late frost, or insect outbreaks (Drobyshev et al., 2014; Hacket-Pain et al., 2015; Hadad et al., 2021). In beech, for instance, drought can produce similar reductions in tree ring width as masting (Ding et al., 2017; Piovesan et al., 2008; Scharnweber et al., 2011). However, since masting is primarily driven by internal carbon and nutrient dynamics (Han et al., 2011; Han and Kabeya, 2017; Pearse et al., 2016), its physiological footprint may extend beyond radial growth alone. Quantitative wood anatomy measures anatomical traits at an intra-annual level, providing insights into tree potential hydraulic conductance (e.g., lumen area, hydraulic diameter, specific conductivity, vessel density), and in relation to carbon and nutrient storage (e.g., tree ring width, earlywood and latewood width, cell wall thickness, degree of lignification, axial parenchyma cells). For these reasons, specific anatomical traits may carry distinct signatures of reproductive investment (Redmond et al., 2019), potentially allowing for a more reliable identification of mast years and opening new possibilities for high-resolution, anatomically based masting reconstructions.

To evaluate the potential of quantitative wood anatomy, we tested the response of ring width and a suite of wood anatomical traits to masting and drought aiming to disentangle the effects of both events. Specifically, we hypothesize that quantitative wood anatomy could help in the identification of a specific “masting signature”, where masting exerts a clearer effect on the traits related to growth and nutrient mobilization, rather than traits involved in plant hydraulics. This analysis laid the basis for the reconstruction of past masting events, and a deeper understanding of the allocation trade-offs associated with masting (Redmond et al., 2019). By successfully linking specific anatomical signatures to masting, our approach represents a significant step forward in understanding long-term reproductive strategies in trees and their interactions with environmental variability.

## Material and Methods

### Sampling site

Trees were sampled in the forest of Carpin in the North-East of Germany (Fig. **1**). The site belongs to the Müritz National Park, which extends over the Mecklenburg Lake District. Specifically, the sampling site (53.20 N; 13.14 E) is in proximity to the Schweingartensee lake, at an elevation of 80 meters above sea level.

**Figure 1.**
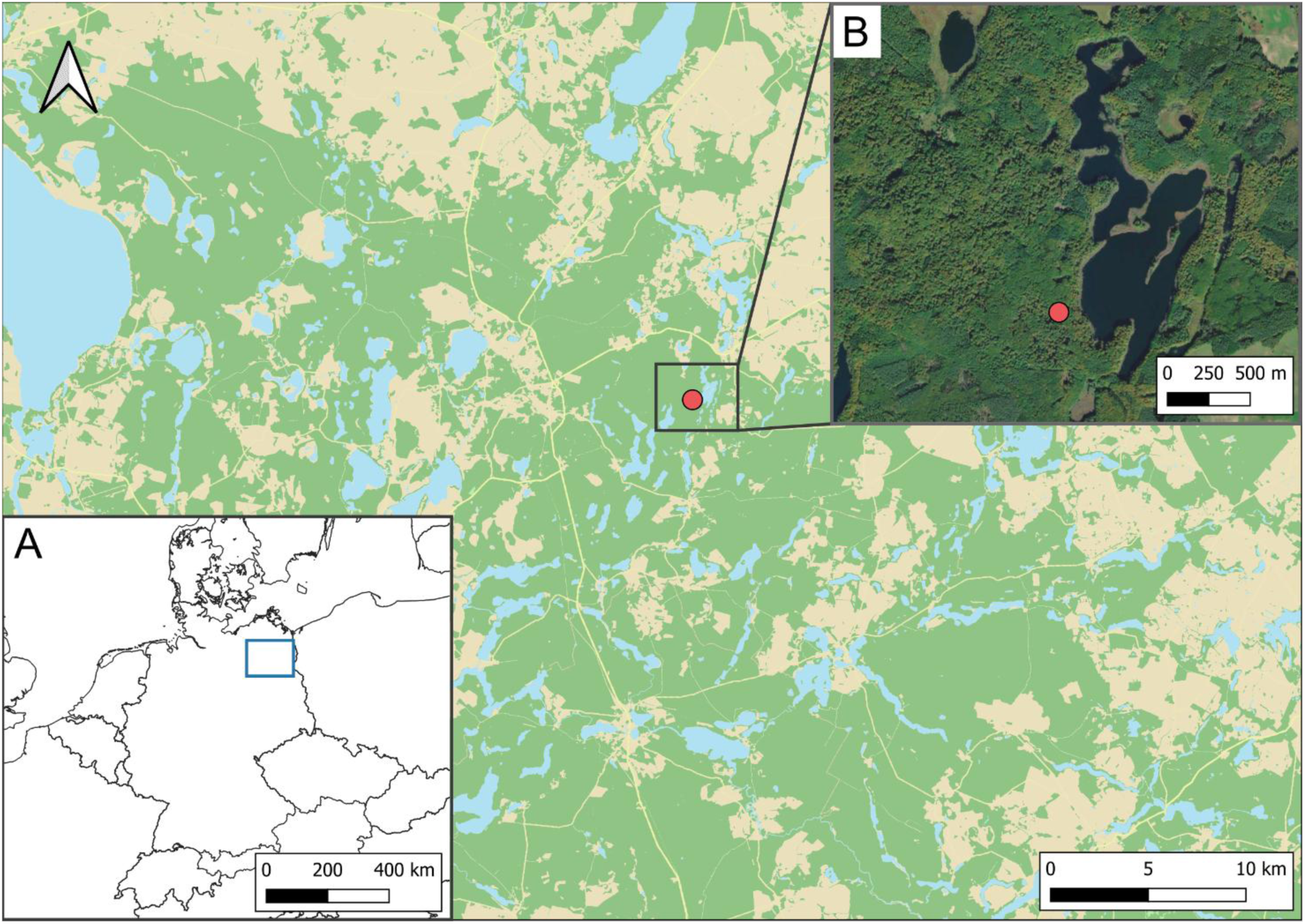
Geographic location of the study site. **(A)** Map of Germany with a subset of the Mecklenburg Lake District (blue rectangle). (**B**) Red dots represent the sampling site located in the Serrahner Wald, close to the Schweingartensee, one of the lakes of the area.

### Sample collection and processing

In February 2022, we sampled twelve healthy, dominant, and fecund trees, collecting two cores per tree at breast height with a 5 mm increment borer. For every tree, GPS position, diameter at breast height, and height measurements were collected. Stem diameter ranged between 62 and 127 centimeters, tree height between 34 and 43 meters.

Cross-dating was performed using one core per tree producing individual tree ring width time series. In the laboratory, cores were mounted on a wooden support, sanded to highlight tree ring visibility, and then scanned with an A3 optical flat-bed scanner with a resolution of 2400 dpi. Tree ring width was then measured with CooRecorder, and crossdated with cDendro 9.3 version (Cybis Electronic 2013). Tree age, together with the collected field data, is reported in Table **S1**. The second core was used for wood anatomical analyses. Cores were cut into 3-4 cm sections and processed with a Leica RM 2245 rotary microtome to produce thin sections of 12 µm thickness. These were stained with a 1:1 Safranin and Astra Blue solution to highlight the lignin and cellulose content, and to improve contrast. Thin sections were mounted on a glass slide and sealed with Euparal mounting media and dried at 65–70 °C for 48 h. Images of the slides were captured using a Zeiss Axio Scan.Z1 slide scanner (Carl Zeiss AG, Germany) with a 10x lens and a resolution of 2.267574 pixel/µm.

### Wood anatomical traits time series

Wood anatomical traits for each individual tree were measured for the period 1970-2021 with the CARROT software (Resente et al., 2024, 2021) on the thin section images. CARROT contains models for measuring tree ring width (TRW), tree ring area computation, and a pretrained model for beech vessel identification. From these analyses, we derived the following traits: coefficient of variation of the lumen area within each ring (CV LA), average hydraulic diameter (Dh), hydraulic diameters of the biggest and the smallest decile of the distribution (Dh min, Dh max), specific conductivity (Ks), and vessel density (VD). Furthermore, CARROT allowed for the creation of a specific model to detect and analyze axial parenchyma cells through the inbuilt retraining function (available at https://zenodo.org/uploads/15262231). This new model allowed the computation of parenchyma cells lumen area (PC LA), number (PC N), and density (PC D). A summary of the abbreviation is provided in Table **1**.

**Table 1.** Table showing traits name and abbreviation. Traits were analyzed within each growth ring, after applying different detrending methods, which resulted in dimensionless measures.

| Abbreviation | Trait |
| --- | --- |
| TRW | Tree ring width |
| CV LA | Coefficient of variation of the lumen area |
| Dh | Hydraulic diameter |
| Dh<sub>min, max</sub> | Hydraulic diameter of the smallest and the biggest decile |
| Ks | Specific conductivity |
| VD | Vessel density |
| PC LA | Lumen area of the parenchymal cells |
| PC N | Number of the parenchymal cells |
| PC D | Parenchyma cells density |
| AWD | Anatomical Wood Density |
| RI | Red Intensity |

We accounted for the vessel widening effect from the pith to the bark due to increasing tree height and crown size (Anfodillo et al., 2013; Lechthaler et al., 2019) as well as the age effect, detrending all the calculated traits with a cubic smoothing spline function with a 50% frequency and a cut-off of 10 years. The optimal cut-off time window was established by selecting the highest R-bar measure among each trait time series (Supporting Information Fig. **S1**). For these analyses we used the detrend function of the dplR Package (Bunn, 2010) that calculates the ratio between the observed and fitted values for each annual tree ring to obtain detrended series (Cook and Kairiukstis, 1990). The function then averages the detrended series employing a bi-weight robust mean to build the mean anatomical chronologies. To account for the ecological phenomenon of masting—which is driven by mechanisms developing over multiple years—and to specifically capture the relevance of interannual temperature variation (i.e., the temperature difference from the previous year) (Vacchiano et al., 2017), we applied a first-order differencing method to detrend the wood anatomical series.

In addition to the wood anatomical traits, we used the methodology described in Resente et al. (in review) to compute the Red Intensity (RI) profile of each tree. RI quantifies the red component in thin sections resulting from the staining process with Safranin and is a proxy for the lignin content in each growth ring averaged by year in order to obtain one data point per year of the chronology. RI was normalized using the slide identification number as a grouping factor and dividing each tree ring RI for the average RI value of the slide. This allowed to avoid bias created by the single thin section thickness. Anatomical Wood Density (AWD) was calculated with the methodology proposed in Kollmann and Côté, (1968) where the percentage of white pixels corresponding to cell lumina was divided by 100 and multiplied by the density of the cell walls (1.53 g/cm³).

### Masting and environmental data

A regional masting chronology was developed using seed production time series retrieved from three different sources: version 2.0 of the MASTREE+ dataset (Hacket-Pain et al., 2022), the publication from Vacchiano et al., (2017), and the German National Inventory FGRDEU (https://fgrdeu.genres.de/erntehandel/ernteaufkommen). Eighteen time series of seed production were selected within a 310 km radius of the sampling site, excluding data from Denmark and Sweden due to their low resolution and discontinuities. This radius was specifically chosen to include the centroids of all the regions adjacent to the study area. To ensure comparability, continuous seed production data were converted into ordinal data (1-5), with 1 representing low and 5 high seed production (Ascoli et al., 2017). Since masting in beech is known to exhibit strong spatial synchrony up to a 300km radius (r∼0.5) (Bogdziewicz et al., 2023; Gallego Zamorano et al., 2018), we first evaluated the relationship between time series similarity and geographic distance from the sampling site using a Mantel test (r² = -0.154, p < 0.05). Recognizing that spatial synchrony tends to decline with increasing distance, we incorporated a distance-based weighting scheme to adjust the influence of each time series, applying the formula: *Correction factor = 1−(Distance/1000)* to each ordinal time series (Table **S2**). In order to obtain a binary classification of masting events (hereafter referred to as masting _Y/N_), where M_Y_ denotes a masting event and M_N_ represents a non-masting event, we applied a threshold criterion to each year for each time series. A given year was classified as M_Y_ if at least one-third of the time series exhibited a value of either 4 or 5. This method led to the identification of 13 masting events (Table 1) from which we excluded 1973, as masting events are unlikely to occur in two successive years. Overall, this was considered as the most reliable evaluation approach since many individual series were incomplete and applying different thresholds than the 1/3 rule would result in an insufficient and non-biologically plausible number of masting events.

To quantify drought, we calculated the Standardized Precipitation Evapotranspiration Index (SPEI) from the temperature and precipitation using the SPEI package (Beguería and Vicente-Serrano, 2023). Climate data were downloaded from the DWD - National Meteorological Service of Germany that holds records of local weather stations. Climate data were sourced from five local weather stations near the sampling site (Müritz, Angermünde, Wittstock, Carpin-Serrahn, and Wesenberg) and averaged to produce complete chronologies for both temperature and precipitation. According to the classification of the SPEI (McKee et al., 1993) and the site-specific conditions, we considered drought events those years for which SPEI is classified as “severely dry” (SPEI _MAY-JUL_ < -1.49). Where SPEI was calculated as the average of the months May, June, and July to account for the importance of spring moisture condition for leaf unfolding (Skvareninova et al., 2024) and for the relevance of early summer drought condition in determining growth reduction (Arnič et al., 2021). As for the masting events, this classification created a binary drought indicator: D_Y_ for drought years and D_N_ for non-drought years (Table **2**).

**Table 2.**
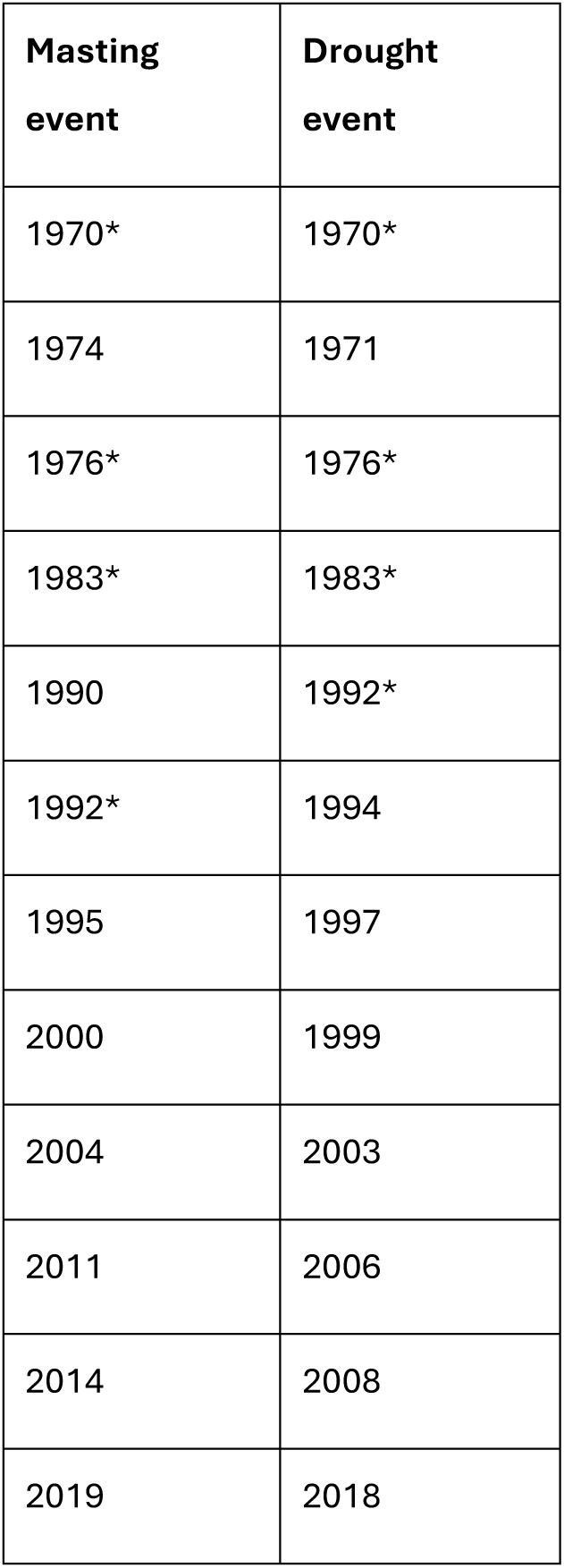
List of the masting and drought events identified from 1970 to 2021. Data were calculated according to local masting records and weather stations. A star was placed for every year in common with both events.

| Masting event | Drought event |
| --- | --- |
| 1970* | 1970* |
| 1974 | 1971 |
| 1976* | 1976* |
| 1983* | 1983* |
| 1990 | 1992* |
| 1992* | 1994 |
| 1995 | 1997 |
| 2000 | 1999 |
| 2004 | 2003 |
| 2011 | 2006 |
| 2014 | 2008 |
| 2019 | 2018 |

### Analysis

To investigate the link between masting and wood anatomy, we first analyzed the main effect of masting on beech TRW and verified that our findings matched results from literature (Drobyshev et al., 2014; Hacket-Pain et al., 2015). Notably, masting in beech involves multiple phenological stages starting from 2 years before flowering and fruiting (t-2) when resource priming occurs, to 1 year before (t-1) when floral buds are initiated, to 1 year after (t+1) where resources remain depleted (Vacchiano et al., 2017). For this reason, we defined the variable “masting phase”, where TRW for each tree and each year was associated to the relative time lag with respect to the year of masting occurrence: TRW _t-2_, TRW _t-1_, TRW _t0_, TRW _t+1_, TRW _t null_, where “t0” is the year of masting and “t null” represents years not associated to any of the defined lag. A Generalized Additive Model (GAM) was used to identify the effect of the masting phases with the following formula (Eq.**1**)

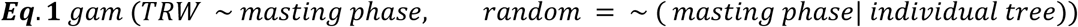

Time interaction was excluded as the TRW time series are detrended. We employed the *estimate<u>_</u> means()* function from the *modelbased* package in R (Hartig et al., 2024) to visualize the mean and the confidence interval of the prediction according to the category and *estimate_contrasts()* function to perform the marginal contrast analysis. P-values were adjusted for multiple testing via the Benjamini-Yekutieli method.

Next, we evaluated the joint and individual effects of masting and drought on wood anatomical traits to determine whether anatomical responses could be attributed to masting alone or confounded by drought. Binary classifications were used for both phenomena (M_Y/N_ and D_Y/N_). Detrended wood anatomical traits were modelled fitting a generalized linear mixed model (GLMM) using Template Model Builder (TMB) according to the following variables: M_Y/N_, D_Y/N_, the cumulative precipitation of the growing season (from March to September) and the mean of the temperature for the same period (Eq. **2**).

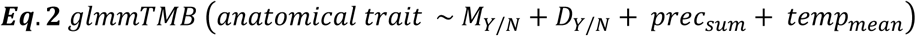

The glmmTMB package allowed for the use of Student’s t distribution which was observed to reach a better fit than normal gaussian for all models except the one involving TRW detrended with the first order difference. Random effect of the single tree was avoided as variance associated to the single trees resulted negligible. Diagnostic tests were performed on all models, including a check on the posterior predictions, collinearity, and uniformity of the residuals using the *performance* package (Lüdecke et al., 2021), and heteroskedasticity, outliers, and overdispersion using the *DHARMa* package (Hartig et al., 2024) (Table **S3**). Differences among drought and masting categories were tested with marginal contrast analysis, and P-values were adjusted for multiple testing via Benjamini-Yekutieli method. Concerning the analysis on the effects of masting and drought, we relied on a confusion matrix to organize the events occurrence, where we defined M_N_D_N_ as the case when neither masting nor drought occurred (the control group), M_Y_D_N_ and M_N_D_Y_ were masting and drought events occurring in isolation, respectively, and M_Y_D_Y_ corresponded to both events occurring in the same year (Table **3**). The comparison between masting and drought effects against none of the events happening (M_Y_D_N_ and M_N_D_Y_ vs M_N_D_N_) constitutes the ground level of our analysis, which was further implemented with the comparison of the effect of M_Y_D_Y_. Results were evaluated with the computation of marginal means from the *model based* package (Makowski et al., 2025) to visualize the mean and the confidence interval of the prediction according to the category, while the *estimate contrast* function was used to perform the marginal contrast analysis. P-values were adjusted for multiple testing via the Benjamini-Yekutieli method.

**Table 3.** Confusion matrix to interpret drought and masting event occurrence. M_N_D_N_ (no masting, no drought), M_Y_D_N_ (masting only) and M_N_D_Y_ (drought only), M_Y_D_Y_ (both masting and drought).

|  | Masting<sub>N</sub> | Masting<sub>Y</sub> |
| --- | --- | --- |
| Drought<sub>N</sub> | $M_N D_N$ | $M_Y D_N$ |
| Drought<sub>Y</sub> | $M_N D_Y$ | $M_Y D_Y$ |

Since the analysis revealed key insights into the effects of masting on wood anatomical traits, we first evaluated trait collinearity using a correlation matrix, highlighting values over 0.7 (Supporting Information Fig. **S2**). Trait selection was then performed with the goal of maximizing the detectable influence of masting on anatomical variation. Specifically, we developed a random forest classification model to predict masting events using three trait groups: 1. QWA: only quantitative wood anatomical traits; 2. TRW: only tree ring width (cubic spline detrended); and 3. QWA + TRW: combined anatomical traits and TRW. Wood anatomical traits considered for the analysis were: RI and VD detrended with the first difference method, and AWD, Dh max, PC N detrended with the cubic spline method. Due to collinearity with TRW, PC N was excluded from the combined model (QWA + TRW).

To assess model performance while accounting for inter-individual structure, we implemented a nested resampling strategy. The outer loop consisted of 12 resamples using group-wise initial splits, with tree identification code as the grouping factor, retaining 75% of the data for training and 25% for testing in each split.

The random forest model was then specified using the *parsnip* package with engine set to *ranger*, and hyperparameters (mtry, min_n, and trees) tuned via grid search (100 combinations). Within each outer resample, we performed inner resampling using 4-fold grouped cross-validation for model tuning. The best model for each split was selected based on the best result of the ROC AUC parameter.

Model evaluation on the outer test sets included the computation of ROC AUC, precision, recall, and F1 score. Variable importance was extracted using impurity-based measures. Additionally, we assessed model interpretability using Individual Conditional Expectation (ICE) plots via the *iml* package (Casalicchio et al., 2025). These plots were generated for each predictor to visualize feature effects on the predicted probability of the positive class.

The same random forest model settings and variable selection were successively applied to predict drought events and understand the performance of the same model on a different target event. ICE plots for drought classification were compared to those for masting classification to understand the extent and nature of each variable’s contribution to the correct prediction.

## Results

### Assessing tree ring width effect

Mean contrast analysis revealed that tree-ring width (TRW) was significantly narrower during mast years (t0) compared to years preceding or following masting events (p < 0.001; Fig. **2**). Among the other time lags, most comparisons were non-significant, with two exceptions: TRW in t–1 was significantly greater than in t–2 (p < 0.05) and t+1 (p < 0.001). (Full p-value results are provided in the supplementary, Table **S4**).

**Figure 2.**
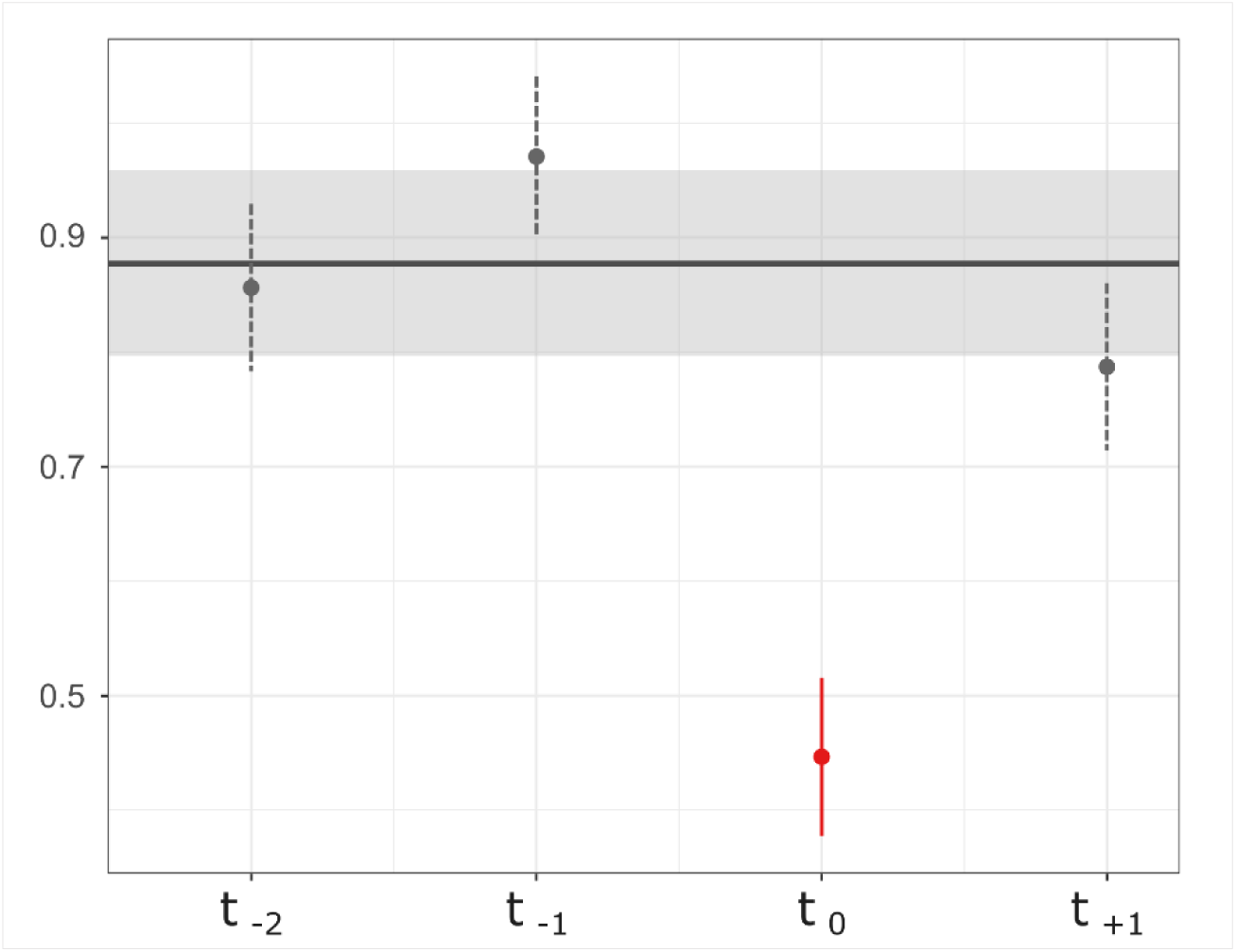
Estimated marginal means on the detrended TRW according to the time lag with respect to the mast year. Confidence intervals set to 95%. Lag categories are represented by t0 year of the masting, t-1 one year prior to the masting, t-2 two years prior to masting, and t+1 one year after masting. Horizontal black line and grey shaded area correspond to the marginal mean and confidence interval (95%) of t null.

### Drought and masting events comparison evaluation

The marginal effects of drought and masting, and their interactions were evaluated across wood anatomical traits. Results are presented by trait groups: 1. ring structural traits (TRW, AWD, and RI), 2. hydraulic traits (Dh, Dh max, Dh min, Ks, VD, and CV LA), and 3. storage and nutrient mobilizations traits (PC LA, PC N, and PC D). Here we show results on the trait detrended with the cubic spline method (Fig. **3**). A table with p-value significance for each comparison is included in the supplementary material (Table **S5**).

**Figure 3.**
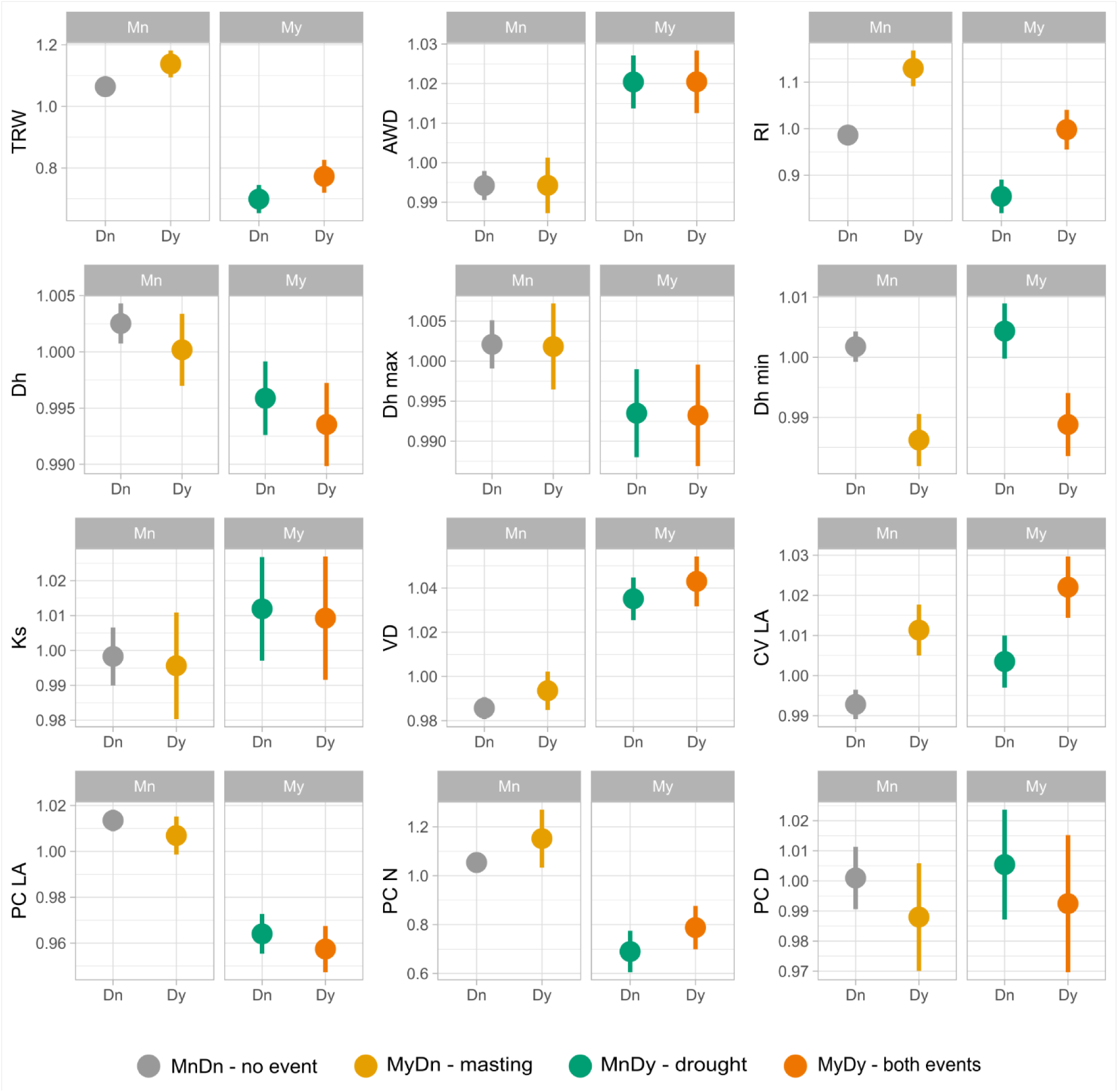
Estimated marginal means of masting events vs drought events for each parameter considered. Time series were detrended with cubic smooth spline with 10 year cutoff. Grey color refers to MnDn (control group), yellow to MnDy (drought), green to MyDn (masting) and orange to MyDy when both events occur.

#### Ring Structural Traits

The group exhibited significant responses across treatment conditions. Tree ring width (TRW) was significantly reduced in mast years compared to non-mast years, whether masting occurred in the presence (MyDy) or absence (MyDn) of drought (p < 0.01). The lowest growth was associated with the coincidence of masting and drought (MyDy). TRW was highest under MnDy, and significantly reduced under MnDn (p <0.01) (Fig. **3**, TRW). Anatomical wood density (AWD) increased significantly under masting, regardless of drought (MyDn, MyDy) (p <0.01), compared to MnDn and MyDy, for which there was no significant difference (Fig. **3**, AWD). Red intensity (RI) did not differ between MnDn and MyDy but was significantly higher under MnDy and significantly lower under MyDn, showing the divergence between anatomy responses when the two single events occur (Fig. **3**, RI).

#### Hydraulic traits

Hydraulic traits showed variable responses to drought, masting, and their combination. Hydraulic diameter (Dh) showed significant reduction in the presence of masting. Dh was lower when masting occurred with respect to non-masting, in the absence of drought (MyDn < MnDn, p <0.01), and the same reduction was observed for both events occurring (MyDy, p <0.01) (Fig. **3**, Dh). Dh max showed the same results, only with a lower degree of significance (p < 0.05). Minimum hydraulic diameter (Dh min) was the only parameter that clearly showed sensitivity only to drought (Fig. **3**, Dh max). Dh min was significantly reduced in drought years (p < 0.01), but there was no detected effect of masting (Fig. **3**, Dh min). Specific hydraulic conductivity (Ks) showed no significant comparison. Vessel density (VD) increased significantly under MyDn and MyDy (p <0.01) with respect to both MnDn and MnDy, which were not significantly different from each other (Fig. **3**, VD). Coefficient of variation of vessel lumen area (CV LA) increased significantly under the combined effects of masting and drought (MyDy), while there was no significant difference among MyDn and MnDy. These two events differed significantly from the MnDn condition (p <0.05 and p <0.01, respectively) (Fig. **3**, CV LA).

#### Storage and Nutrient Mobilization Traits

These traits were also significantly affected by masting, and to a lesser degree by drought. Parenchyma cell lumen area (PC LA) was significantly lower in MyDn and MyDy with respect to both MnDn and MnDy (p <0.01) (Fig. **3**, PC LA). Parenchyma cell number (PC N) followed the same pattern as PC LA but, while for PC LA masting and drought seemed to have the same effect that was significant only for masting, in PC N masting and drought exhibit opposite behavior with respect to the control (MnDn) (Fig. **3**, PC N). However, the confidence interval revealed that masting is the only significant event affecting this trait. Parenchyma cell density (PC D) showed no significant comparisons.

Repeating the analysis for the same traits detrended with the first order difference method produced largely similar results, with differences only for certain traits. Specifically, within the ring structural traits, comparison between MnDy and MnDn in TRW were no longer significant, while AWD MnDy versus MnDn showed significant difference, with higher values for MnDy, and even greater difference for MyDn. RI relationship remained unchanged, aside from the MnDn vs MyDy comparison which differed significantly. For the hydraulic traits, the only notable difference regarded CV LA, with a heightened difference between MyDn and MnDy, where masting showed no significant difference with MnDn, as well as drought compared to MyDy. Similarly to the other groups of traits, for the storage and nutrient mobilizations traits only PC LA exhibited difference due to the detrending method and specifically heightened the difference between MnDy towards MnDn, and MyDn towards MyDy. Results for the traits detrended with the first order difference method are available in the supplementary material (Supporting Information Fig. **S3**) along with the table with p-value significance for each comparison (Table **S5**).

#### Results summary

Masting, drought and their interaction significantly influenced wood anatomical traits, which were analyzed grouped into structural, hydraulic, and storage-related categories. Analyses based on cubic spline detrended data revealed distinct patterns:

Ring Structural Traits: Tree-ring width (TRW) was highest during drought-only years and lowest when both drought and masting co-occurred. Anatomical wood density (AWD) increased with masting, while red intensity (RI) responded in opposite directions under drought and masting alone.

Hydraulic Traits: Masting led to narrower vessel diameters (Dh, Dh max), while Dh min was the only tested trait that showed responses to drought but not to masting. Vessel density (VD) rose with masting, and lumen area variation (CV LA) increased when both events occurred. Specific hydraulic conductivity (Ks) showed no clear response.

Storage and Nutrient Traits: Masting caused a marked reduction in parenchyma cell lumen area (PC LA) and number (PC N), suggesting a trade-off in storage capacity. Cell density (PC D) remained unaffected.

### Mast year prediction

The predictive performance analysis revealed that the combined QWA+TRW model achieved marginally higher metrics (F1-score: 0.91±0.006; precision: 0.88±0.01; recall: 0.95±0.02; ROC AUC: 0.86±0.02) compared to the QWA-only model (F1-score: 0.91±0.01; precision: 0.87±0.01; recall: 0.92±0.02; ROC AUC: 0.85±0.03). However, the results varied across runs and showed overlapping confidence intervals between QWA+TRW and QWA-only model. In contrast, the TRW-only model consistently demonstrated the lowest performance across all metrics (F1-score: 0.88±0.01; precision: 0.84±0.01; recall: 0.92±0.05; ROC AUC: 0.79±0.02). The narrow error bars observed for most metrics (Fig. 4a) indicated stable model performance across resampling iterations.

**Figure 4.**
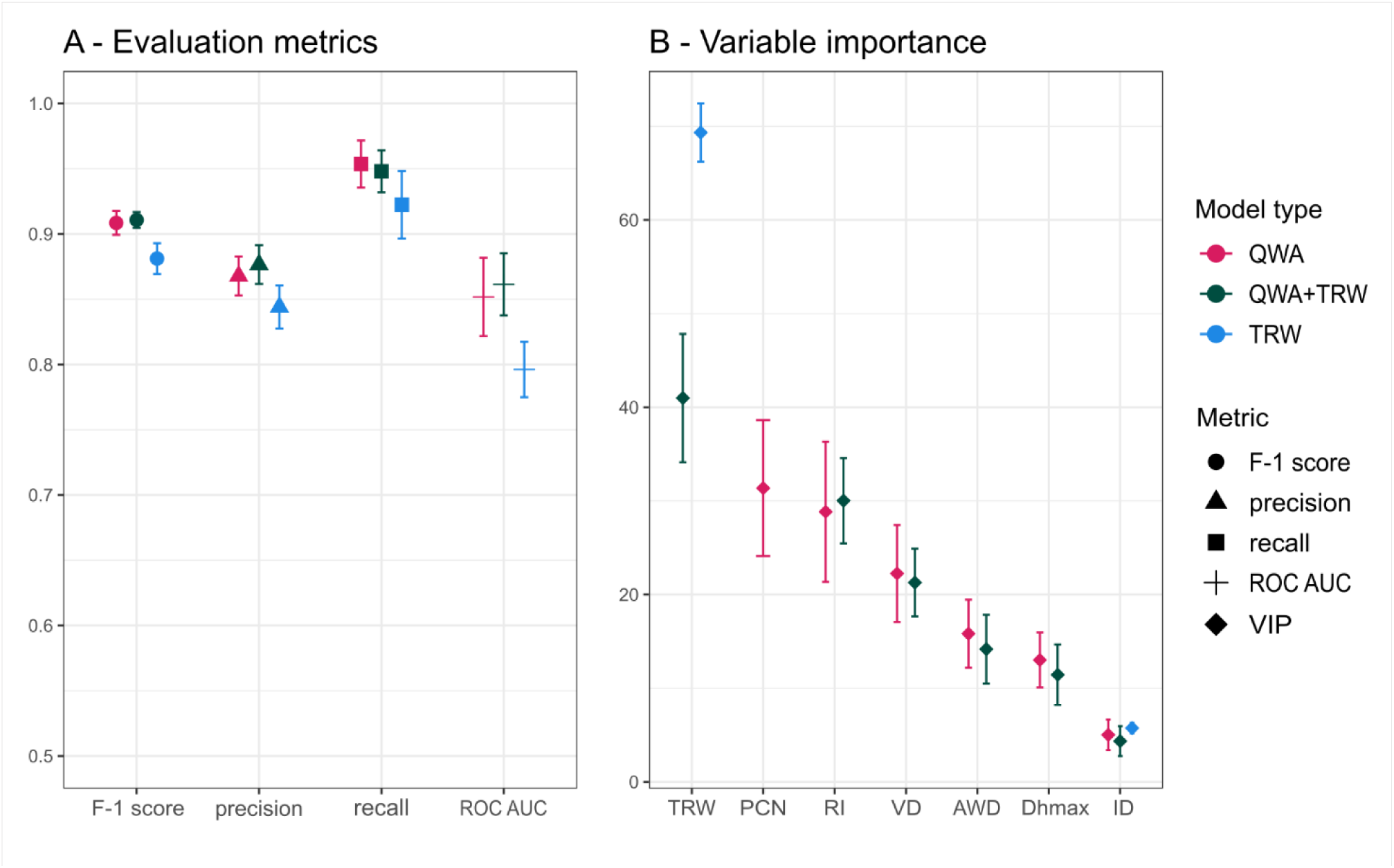
Performance evaluation and variable importance of Random Forest models in predicting masting behavior. *(A)* Evaluation metrics (F-1 score, precision, recall, and ROC AUC) for three model types: QWA (quantitative wood anatomical traits only), QWA + TRW (combined wood anatomical traits and tree-ring width), and TRW (tree-ring width only). Standard deviation bars indicate variability across cross-validation folds. *(B)* Variable importance rankings (VIP) of the top predictors in the Random Forest models. Error bars represent variability in importance estimates across models. Traits are labeled according to the wood anatomical traits considered; all traits were detrended using the cubic spline method, except for RI and VD, which were detrended using the first order difference method.

Mean variable importance for the combined model (QWA+TRW) showed that the most influential predictor was TRW, followed by RI, and VD. Other traits such as AWD and Dh max also contributed to the model but with comparatively lower importance. The sample ID ranked lowest in importance, indicating minimal influence from individual-specific variation. Beside this difference the two groups perform similarly in the variable importance evaluation (Fig. 4b).

Results on the same models applied to predict drought events were also assessed and showed that the main evaluation parameter – ROC AUC – was substantially lower for the same three categories of traits (TRW = 0.54, QWA = 0.68, and TRW + QWA = 0.68). Recall and precision showed more divergent results, with higher recall for all three groups, but lower precision (Supporting Information Fig. **S4**).

In this regard, ICE plots were evaluated to assess the nature of the prediction of masting and drought according to the same QWA variables. Results showed that VD and AWD show a positive relationship with masting probability: as values increase, so does the likelihood of a masting event. RI and PCN exhibit a strong negative relationship: lower values of these traits are associated with higher probabilities of masting. The same is verified for Dh max but with lower intensity. Tree identification code (ID) shows relatively minor variation, indicating that individual-level effects are less critical than anatomical traits for predicting masting (Fig. **5**). The same traits were evaluated for drought prediction, where we found ICE curves to be generally flatter and more diffuse, indicating weaker predictive relationships for drought of these traits. RI together with AWD are the only traits showing a moderate inflection at intermediate values. Dh max, AWD, PC N, and ID show only subtle changes across their range, supporting the earlier observation that drought events do not induce strong or consistent anatomical changes.

**Figure 5.**
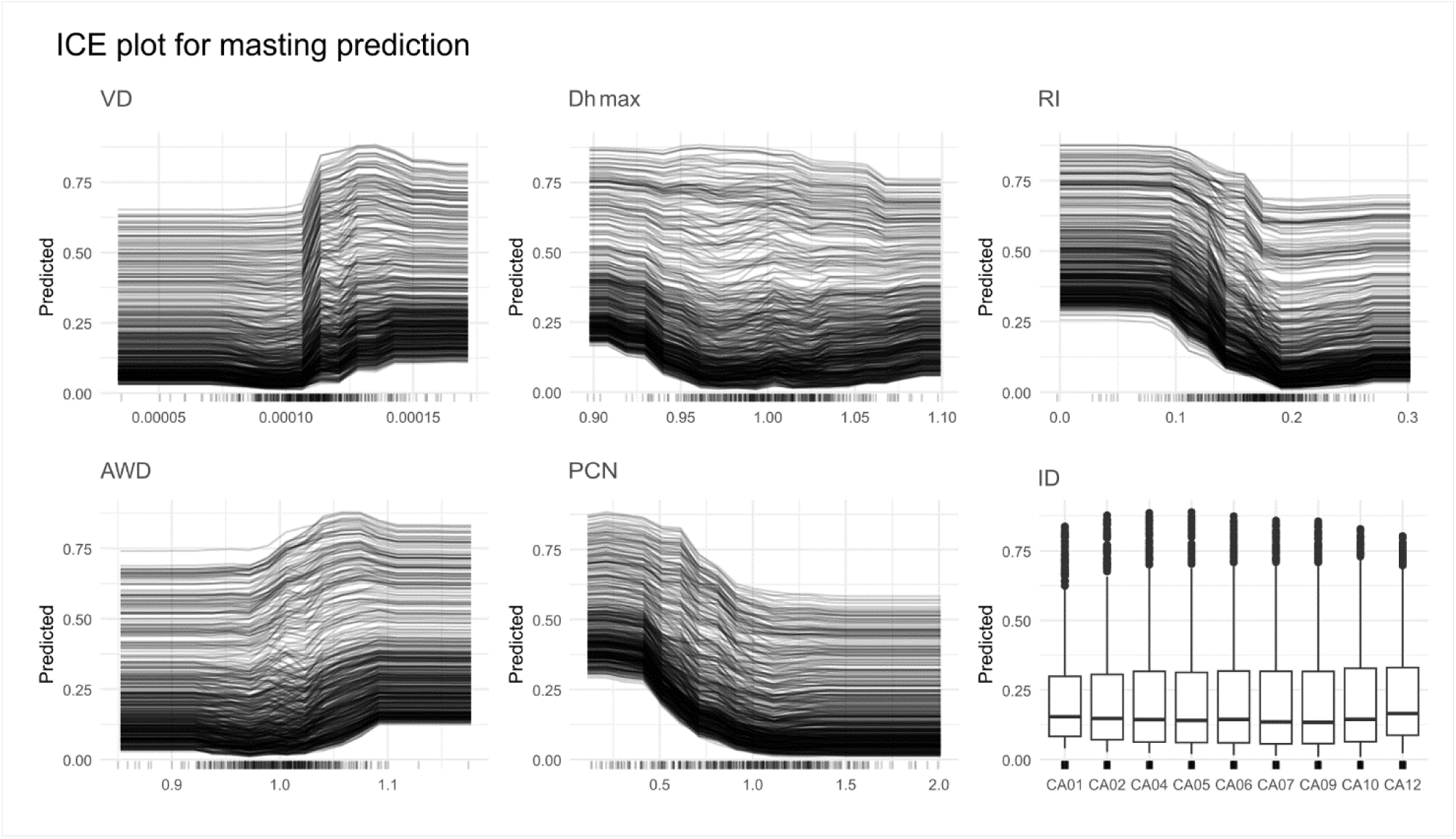
Individual Conditional Expectation (ICE) plots for masting prediction models. Each line represents an individual observation, illustrating how predicted probability changes across the range of each variable while all other variables are held constant. ICE plots for masting prediction showing the influence of vessel density (VD), maximum hydraulic diameter (Dhmax), red intensity (RI), anatomical wood density (AWD), number of parenchyma cells (PC N), and tree identification code (ID) on the model’s predicted probability of a masting year

## Discussion

Our results demonstrated that wood anatomical traits contain a clear and consistent imprint signal of masting events, and that the responses are distinct from those induced by drought. Results enabled us to reliably predict the occurrence of large mast years in beech avoiding confounding signals like drought. These findings suggest a high potential for masting reconstructions based on quantitative wood anatomy (QWA) representing a significant advancement compared to methods based on tree ring width (TRW) (Hacket-Pain et al., 2019).

### Effects of masting and drought on wood anatomy

The distinct anatomical signatures of masting and drought observed in our study provide mechanistic insights into how trees balance resource allocation under different stressors.

We verified that our findings are in line with literature in terms of the effect of masting on TRW, which, predominantly in beech but also in other species, was reported to cause ring width reduction (Drobyshev et al., 2014; Hacket-Pain et al., 2024, 2015; Selås et al., 2002). Our analysis further indicates that this reduction is evident not only when comparing masting to non-masting years but also when considering the physiological patterns preceding or following beech masting events, such as floral induction, resource priming the previous year, and resource depletion the year after masting (Vacchiano et al., 2017) (Fig. **2**).

While it is well established that beech masting reduces growth, the effects of masting on wood anatomical traits alone and in combination to drought remain largely unexplored, with only one study specifically tackling this research gap, investigating the effects of masting on beech hydraulic structure (Rodríguez-Ramírez et al., 2019). However, relying on macroscopical images of beech cores holds the disadvantage of poorly capturing vessel variability. Disentangling drought from masting effects was essential to isolate the true effect of reproductive investment on quantitative wood anatomy, to reveal distinct physiological trade-offs that would otherwise be obscured by overlapping stress responses. A general observation on the effect of drought and masting on wood anatomy, is that drought did not seem to have a strong impact on our samples. This is primarily supported by the fact that TRW was not reduced by the occurrence of drought, as it is otherwise widely documented in beech (Leuschner et al., 2001; Martinez del Castillo et al., 2022; Schmied et al., 2023). Generally, the main effect of drought was observed in hydraulic traits, with the hydraulic diameter of the smallest cells (Dh min) being clearly negatively affected solely by drought. These findings align with well-established physiological responses to water limitation, where trees prioritize hydraulic safety by narrowing conduits to reduce embolism risk (Anderegg et al., 2016; Arend et al., 2022). However, the effect of drought on the other hydraulic traits is partially masked by the effect of masting, generating in the mean and max hydraulic diameter (Dh and Dh max), specific hydraulic conductivity (Ks) and coefficient of variation of the lumen area (CV LA) wide error bars and overlapping responses that lead to non-significant comparison between drought and masting effects (Fig **3**). This lack of specific response to drought by a certain array of traits might be explained by the fact that the sampling site is located in the Mecklenburg Lake District, which constitutes a moist environment where microclimate is significantly influenced by lakes (Scharnweber et al., 2011). Moreover, the conspicuous tree height (34 - 42.9 m) also suggests a fertile soil and a generally favorable growing condition.

By studying masting in combination with drought, we were able to disentangle the specific impact of masting on quantitative wood anatomy, providing novel insights into their interaction. This comparison revealed that masting influenced anatomical, while drought alone did not show statistically significant changes compared to the control group (MnDn). The broader anatomical signature of masting is represented by reductions in tree-ring width (TRW), red intensity (RI), mean and maximum hydraulic diameter (Dh and Dh max), lumen area and number of parenchyma cells (PC LA and PC N), coupled with increases in vessel density (VD) and anatomical wood density (AWD). These traits and the nature of their responses suggested a complex mechanism of general downregulation of growth and investment in xylem tissue during masting, while reflecting a compensatory strategy to maintain hydraulic function and mechanical stability under resource-constrained conditions. Evidence suggests, in fact, that ecological mechanisms such as the ones driving masting effects on growth, should not be reduced to simple carbon-mediated ‘trade-off’ (Mund et al., 2020), as they also involve structural adjustments tied to whole-tree hydraulic architecture. Previous work on *Fagus sylvatica* indicates that tree size and crown architecture better explain vessel diameter variation, rather than general climatic factors (Kröner et al., 2024; Prislan et al., 2018). In this context, our findings align with observations that mast years are associated with reduced shoot length and fewer leaves per shoot (Han et al., 2011; Müller-Haubold et al., 2015), pointing towards the link between reduced leaf area and lower conductive demand (decrease in Dh).

Notably, this study is among the first to explore such a broad suite of wood anatomical traits in relation to masting, including a specific focus on axial parenchyma characteristics (Plavcová et al., 2024; Słupianek et al., 2021). Parenchyma cells are primarily involved in nutrient storage and mobilization functions (Tomasella et al., 2020), and while a reduction in their number (PC N) was expected due to narrower tree rings and known trade-offs with hydraulic traits (Zheng and Martínez-Cabrera, 2013), we also found a significant decrease in parenchyma lumen area (PC LA). This finding is particularly interesting considering the existing literature reporting high genetic control and low plasticity of wood anatomical features (Chhetri et al., 2020; Hajek et al., 2016). A plausible explanation is that during mast years, when non-structural carbohydrates (NSC) used for reproduction are predominantly sourced from current-season assimilates (Hoch, 2005; Ichie et al., 2013), there is a reduced reliance on stored reserves. As a result, there may be limited carbon investment into storage tissue development, which further contributes to explain the concurrent reduction in both the number and lumen area of parenchyma cells.

Red intensity (RI), a proxy for lignin content in tree rings, showed opposite effects for masting than for drought (Fig. **3**) (Resente et al., in review). This distinction is ecologically meaningful as lignin supports both structural defense and hydraulic safety (Polo et al., 2020). The RI reduction during masting aligns with reported carbon trade-offs in reproductive years, where lignin biosynthesis may be deprioritized to allocate resources toward flowering and seed production (Han and Kabeya, 2017; Monks and Kelly, 2006; Redmond et al., 2019). Conversely, drought-induced RI increase mirrors global patterns of enhanced lignification under water stress to reinforce xylem against embolism (Crivellaro et al., 2022; Lima et al., 2018; Resente and Crivellaro, 2025). These opposing RI trends empirically validate theoretical models predicting resource allocation shifts between reproduction and stress resilience (Bogdziewicz, 2022).

### The potential of wood anatomy for masting reconstruction

Previous attempts to reconstruct ecological events such as masting have relied solely on tree-ring width (TRW) as anatomical proxy. For these reason existing reconstruction studies, either rely on the combination of TRW and climatic data, or, in the case of Araucaria, employ the dioecious characteristic of the species to enable the separation of masting from the climate signal (Drobyshev et al., 2014; Hacket-Pain et al., 2024; Mundo et al., 2021). Given that masting influences forest regeneration, carbon cycling, and wildlife dynamics (Bogdziewicz et al., 2021), accurate reconstructions are crucial for understanding long-term ecosystem responses to environmental change. Our findings demonstrate that although TRW carries substantial predictive power, it is outperformed by models incorporating quantitative wood anatomical traits (QWA) or models including a combination of QWA and TRW. To our knowledge, this is one of the first studies to establish a direct link between masting events reconstruction and a wide suite of QWA traits, beside Rodríguez-Ramírez et al. (2019) which focused on the hydraulic structure. Particularly, the QWA-only model performed as well as the combined model, confirming that QWA traits contain substantial independent information relevant to masting prediction. Among the QWA traits, the number of parenchyma cells (PC N) and red intensity (RI) emerged as particularly strong predictors (Fig. 4). For PC N, this can be attributed to the expected collinearity between PC N and TRW, since narrower tree rings are generally associated with a lower number of cells (Babushkina et al., 2021). For RI, the opposite effects of drought and masting are relevant; drought increased while masting decreased RI. This provides a strong basis for robustly identifying mast years, which have previously been challenging due to common response of TRW to these two drivers (Drobyshev et al., 2014). The strong predictive responses of both PC N and RI validated our analytical approach to isolating the effects of masting; while traits such as VD, AWD, and Dhmax showed comparatively weaker predictive power, suggesting a less distinctive response to masting events.

When the same random forest modeling approach was applied to drought prediction, model performance dropped significantly across all metrics, particularly in relation to ROC AUC, which indicates the general reliability of the model in classifying the event as drought or non-drought. This suggests that traits selected to maximize the effect of masting on wood anatomy do not capture drought-specific signatures with the same reliability. The moderate performance predicting drought may be largely driven by the discriminative role of RI, which in the previous analysis (Fig. **3**) and in the ICE plots (Fig. **5**) showed a distinct opposite response to drought than to masting. Nonetheless, the generally lower precision and ROC AUC values indicate that anatomical traits selected better reflect the coordinated, resource-driven shifts associated with reproductive events. The ICE plots further support this interpretation. Traits such as vessel density (VD), red intensity (RI), and parenchyma cell number PCN show clear, directional relationships with masting probability, reflecting their strong influence on prediction. In contrast, ICE plots for drought prediction display flatter, more diffuse patterns, indicating that variability in these traits does not translate into consistent predictive signals for drought years. This suggests that while the same anatomical dataset may contain some signal of drought impact as one third of the mast years were also drought years, it lacks the distinct, coordinated anatomical signature seen in response to masting. Further support for the general applicability of our approach comes from the finding that individual tree identity, despite the low sample size, had negligible influence on model performance, suggesting that the predictive framework could be reliably extended across different trees and potentially across sites, which constitutes an important step toward developing scalable, species-wide reconstructions of masting dynamics.

Together, these findings show that QWA traits not only improve predictive accuracy but also provide insight into the physiological adjustments that trees undergo under biotic and abiotic pressures. Moreover, in line with previous dendrochronological reconstructions (Drobyshev et al., 2014; Mundo et al., 2021), our binary mast/non-mast approach lays the groundwork for future studies aiming to reconstruct continuous seed production—a crucial step toward detecting long-term trends and potential masting patterns.

### Conclusions

In this study we present the first clear link between masting, the synchronized and variable seed production, and structural changes in the wood anatomy of European beech.

Our results show that mast years are associated with a general downregulation of growth and storage investment, evident in narrower rings, smaller hydraulic diameters, reduced lignin content, and fewer and smaller parenchyma cells, coupled with increased vessel density and anatomical wood density. These patterns suggest a complex mechanism of carbon reallocation towards reproduction, while reflecting a compensatory strategy to maintain hydraulic function and mechanical stability under resource-constrained conditions. Notably, red intensity – a proxy reflecting lignin content in tree rings – was the only trait to show opposite responses to masting and drought, emphasizing the distinct ecological strategies trees adopt under reproductive versus abiotic stress.

By capturing these specific wood anatomical adjustments, QWA traits emerged as strong predictors of masting. The classification model based on QWA performed better than a model including TRW alone, highlighting the advantage of the intra annual approach, and equally to a combined model of QWA and TRW. Moreover, the most explanatory traits for predicting masting events mirrored in intensity and nature the results found on the effect of masting in wood anatomy. The minimal effect of individual tree identity further supports the possibility of generalization of our approach, promoting species-wide reconstructions based on anatomical data. Importantly, the same anatomical traits that reliably predicted masting, failed to accurately capture drought years in our dataset. This discrepancy further reinforces that the anatomical signature observed is tightly linked to reproductive effort rather than overlapping environmental stress. In addition, the study site presents particularly favorable conditions for tree growth, therefore, we recommend replicating the study in a more drought-sensitive location.

These findings underscore that quantitative wood anatomy provides a robust, species-wide tool for reconstructing and predicting mast years in European beech, bridging the current gap on reliable reconstruction methodologies, and therefore enabling more accurate predictions of future tree reproduction trajectories.

## Acknowledgements

DA and AP were supported by the Agritech National Research Center and the European Union Next-Generation EU (PIANO NAZIONALE DI RIPRESA E RESILIENZA (PNRR) – MISSIONE 4 COMPONENTE 2, INVESTIMENTO 1.4 – D.D. 1032 June 17, 2022, CN00000022), Spoke 4, Task 4.1.3

## Competing interests

The authors have no competing interests to declare.

## Author contributions

All authors contributed equally to the study. Conceptualization: GR, MW, AC, AP, and DA. Design: GR and MW. Data Collection: GR, EF and FM. Analysis: GR, EF, FM, AHP, DA and AP. Original draft: GR, DA, AHP. Review & Editing: GR, AC, EF, AP, FM, MW, AHP, RM, and DA. Supervision: DA, RM, AHP, AP, and AC.

## Data availability

The data that support the findings of this study are available from the corresponding author, GR, upon request.

## Supplementary information legend

Figure S1. R-bar evaluation performed on data from the complete set of traits detrended with four detrending method (cubic spline with a 10 years’ time window, 20 years, 30 years and first order difference)

Figure S2. Correlation matrix between all the traits detrended with both cubic spline 10 years’ time window and first order difference methods.

Figure S3. Estimated marginal means of masting events vs drought events computed for every parameter considered in the analysis detrended with first order difference.

Figure S4. Performance evaluation and variable importance of Random Forest models in predicting drought behavior.

Table S1. Field collected data, together with derived data from dendrochronological analyses.

Table S2. Masting time series (1970-2021) Id number of each column is composed of "site id"_"NUTS1"_"source"

Table S3. Models’ diagnostic tests

Table S4. Estimated marginal means metric with contrast applied to the TRW trait. P value adjusted via the Benjamini-Yekutieli method.

Table S5. P-value results for significance for each comparison (Mn vs My vs Dn vs Dn) for both detrending methods

## Notes

### Competing Interest Statement

The authors have declared no competing interest.

### Summary of Updates

Version uploaded contains only the original manuscript, replacing the previous proof of submission.

